# The promise of disease gene discovery in South Asia

**DOI:** 10.1101/047035

**Authors:** Nathan Nakatsuka, Priya Moorjani, Niraj Rai, Biswanath Sarkar, Arti Tandon, Nick Patterson, Gandham SriLakshmi Bhavani, Katta Mohan Girisha, Mohammed S Mustak, Sudha Srinivasan, Amit Kaushik, Saadi Abdul Vahab, Sujatha M. Jagadeesh, Kapaettu Satyamoorthy, Lalji Singh, David Reich, Kumarasamy Thangaraj

**Author notes:** co-senior authors (David Reich, Kumarasamy Thangaraj).

## Abstract

The more than 1.5 billion people who live in South Asia are correctly viewed not as a single large population, but as many small endogamous groups. We assembled genome-wide data from over 2,800 individuals from over 260 distinct South Asian groups. We identify 81 unique groups, of which 14 have estimated census sizes of more than a million, that descend from founder events more extreme than those in Ashkenazi Jews and Finns, both of which have high rates of recessive disease due to founder events. We identify multiple examples of recessive diseases in South Asia that are the result of such founder events. This study highlights an under-appreciated opportunity for reducing disease burden among South Asians through the discovery of and testing for recessive disease genes.

---

South Asia is a region of extraordinary diversity, containing over 4,600 anthropologically well-defined groups many of which are endogamous communities with significant barriers to gene flow due to cultural practices that restrict marriage between groups^1^. Of the tiny fraction of South Asian groups that have been characterized using genome-wide data, many exhibit large allele frequency differences from close neighbors^2–4^, consistent with strong founder events whereby a small number of ancestors gave rise to many descendants today^4^. The pervasive founder events in South Asia present a potential opportunity for reducing disease burden in South Asia. The promise is highlighted by studies of founder groups of European ancestry – including Ashkenazi Jews, Finns, Amish, Hutterites, Sardinians, and French Canadians – which have resulted in the discovery of dozens of recessive disease causing mutations in each group. Prenatal testing for these mutations has substantially reduced recessive disease burden in all of these communities^5,6^.

We carried out new genotyping of 1,663 samples from 230 endogamous groups in South Asia on the Affymetrix Human Origins single nucleotide polymorphism (SNP) array^7^. We combined the data we newly collected with previously reported data, leading to four datasets (Figure 1a). The Affymetrix Human Origins SNP array data comprised 1,955 individuals from 249 groups in South Asia, to which we added 7 Ashkenazi Jews. The Affymetrix 6.0 SNP array data comprised 383 individuals from 52 groups in South Asia^4,8^. The Illumina SNP array data comprised 188 individuals from 21 groups in South Asia^9^ and 21 Ashkenazi Jews^9,10^. The Illumina Omni SNP array data comprised 367 individuals from 20 groups in South Asia^11^. We merged 1000 Genomes Phase 3 data^12^ (2,504 individuals from 26 different groups including 99 Finns) with each of these datasets. We removed SNPs and individuals with a high proportion of missing genotypes or that were outliers in Principal Components Analysis (PCA) (Figure 1b; Supplementary Text). The total number of unique groups analyzed in this study is 263 after accounting for groups represented in multiple datasets. To our knowledge, this represents the richest set of genome-wide data from anthropologically well-documented groups from any region in the world.

**Figure 1.**
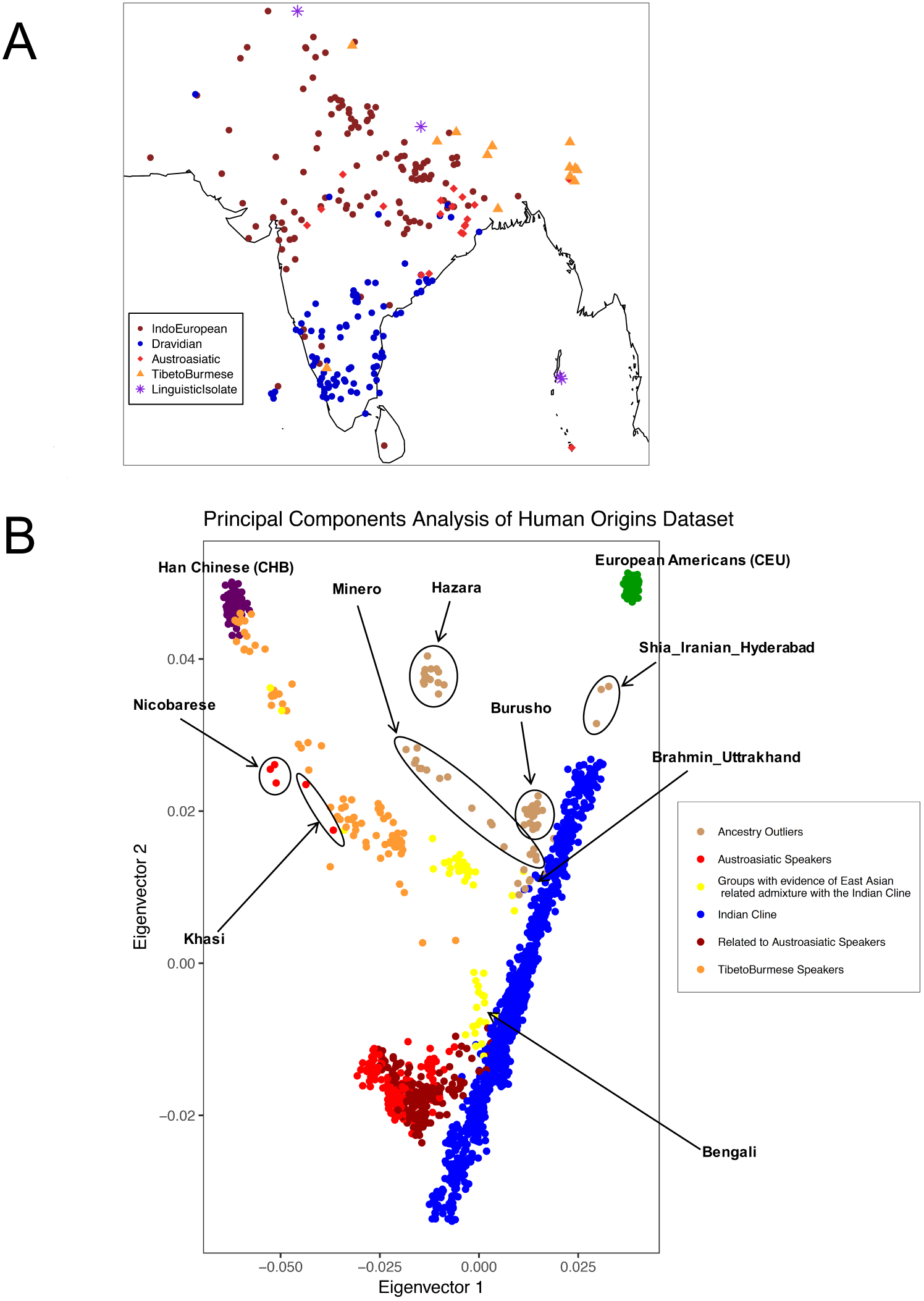
Dataset overview. (a) Sampling locations for all analyzed groups. Each point indicates a distinct group (random jitter was added to help in visualization at locations where there are many groups). (b) PCA of Human Origins dataset along with European Americans (CEU) and Han Chinese (CHB). There is a large cluster (blue) of IndoEuropean and Dravidian speaking groups that stretch out along a line in the plot and that are well-modeled as a mixture of two highly divergent ancestral populations (the “Indian Cline”). There is another larger cluster of Austroasiatic speakers (light red) and groups that cluster with them genetically (dark red). Finally, there are groups with genetic affinity to East Asians that include Tibeto-Burman speakers (orange) and those that speak other languages (yellow).

We devised an algorithm to quantify the strength of the founder events in each group based on Identity-by-Descent (IBD) segments, large stretches of DNA shared from a common founder in the last approximately one hundred generations (Figure 2). We computed an “IBD score” as a measure for the strength of the founder event in each group’s history: the average length of IBD segments between 3-20 centimorgans (cM) shared between two genomes normalized to sample size. Since we are interested in characterizing the impact of recessive diseases that do not owe their origin to consanguineous marriages of close relatives, we ignored self-matches (internal homozygosity) in IBD calculations. We removed all individuals that had evidence of recent relatedness (within a few generations) to others in the dataset by computing IBD between all pairs of individuals in each group and removing one individual from the pairs with outlying numbers of IBD segments (our focus on founder events rather than recent relatedness also explains our choice to exclude IBD segments of greater than 20 cM in size). We validated the effectiveness of this procedure by simulation (Supplementary Table 1; Online Methods).

**Figure 2.**
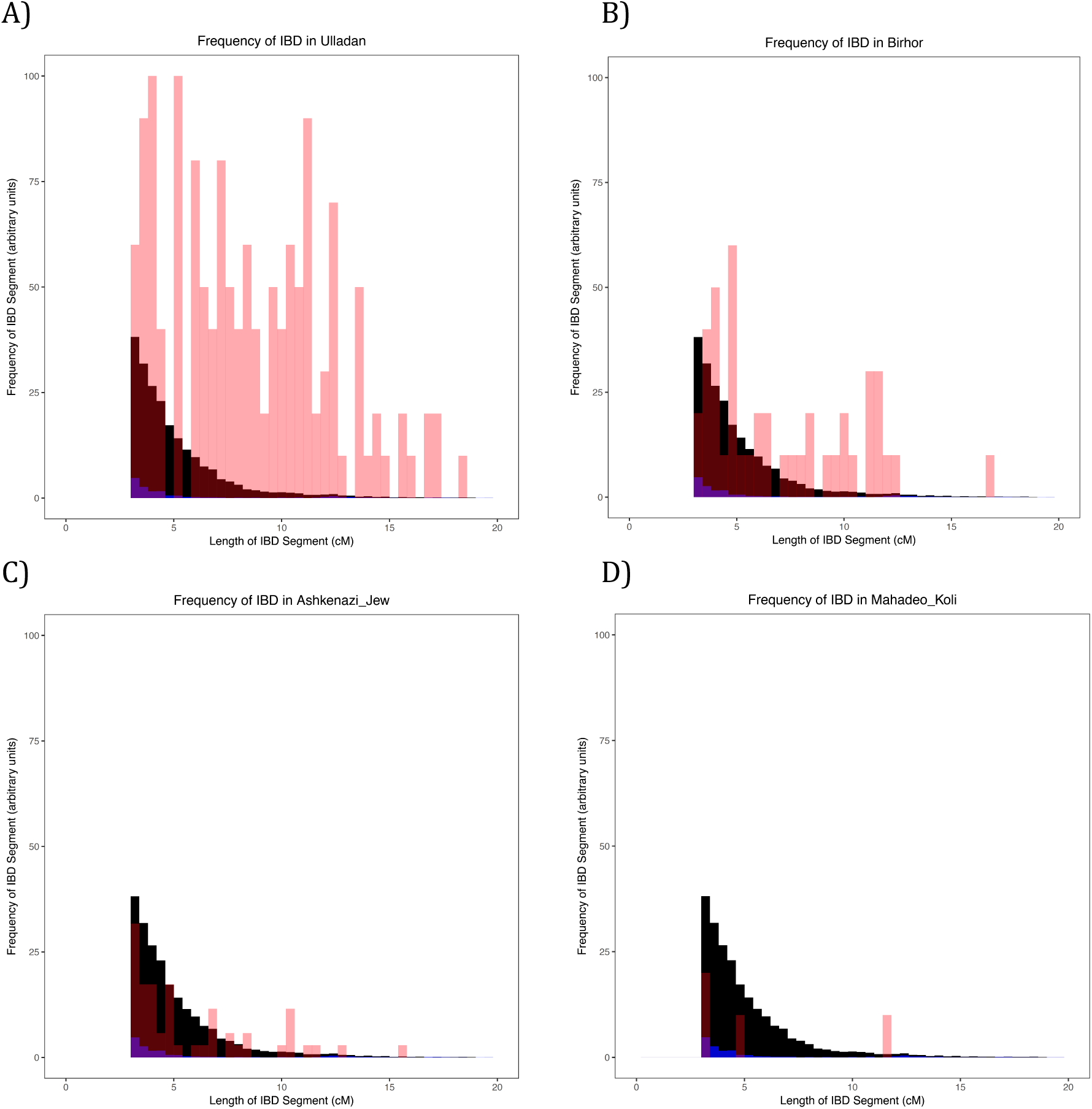
Example histograms of IBD segments to illustrate the differences between groups with founder events of different magnitudes: These histograms provide visual illustrations of differences between groups with different IBD scores. As a ratio relative to Finns (FIN; black), these groups (red) have IBD scores of: (A) ∼26 in Ulladan, (B) ∼3 in Birhor, (C) ∼0.9 in Ashkenazi Jews, and (D) ∼0.1 in Mahadeo_Koli. In each plot, we also show European Americans (CEU) with a negligible founder event in blue. Quantification of these founder events is shown in Figure 3 and Online Table 1. The IBD histograms were normalized for sample size by dividing their frequency by 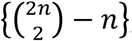, where *n* is the number of individuals in the sample. All data for the figure are based on the Human Origins dataset.

We expressed IBD scores for each group as a fraction of the IBD scores of the 1000 Genomes Project Finns merged into each respective dataset. Due to the fact that all the SNP arrays we analyzed included more SNPs ascertained in Europeans than in South Asians, the sensitivity of our methods to founder events is greater in Europeans than in South Asians, and thus our estimates of founder event strengths in South Asian groups are conservative underestimates relative to that in Europeans (Supplementary Figure 1 demonstrates this effect empirically and shows that it is less of a bias for the strong founder events that are the focus of this study). We computed standard errors for these ratios by a weighted Block Jackknife across chromosomes and declared significance where the 95% confidence intervals did not overlap 1. We carried out computer simulations to validate our procedure. The simulations suggest that we are not substantially overestimating the magnitudes of modest founder events, since for a simulated founder event that is half the magnitude of that in Finns, we never infer the score to be significantly greater than in Finns. The simulations also suggest that our procedure is highly sensitive to detecting strong founder events, since for sample sizes of at least 5, the algorithm’s sensitivity is greater than 95% for determining that a group with two times the bottleneck strength as Finns has an IBD score significantly greater than that of Finns (Supplementary Figure 2 and Supplementary Table 2). We also used two additional non-IBD based methods to measure the strength of founder events and in cases where a comparison was possible found high correlation of these results with our IBD scores (Supplementary Text and Supplementary Table 3).

We infer that 81 out of 263 unique groups (96 out of 327 groups if not considering the overlap of groups among datasets) have an IBD score greater than those of both Finns and Ashkenazi Jews (Figure 3). These results did not change when we added back the outlier samples that we removed in quality control. A total of 14 of these groups have estimated census sizes of over a million (Figure 3; Table 1). However, the groups with smaller census sizes are also very important – outside of South Asia, small census size groups with extremely strong founder events such as Amish, Hutterites, and people of the Saguenay Lac-St. Jean region have led to the discovery of dozens of novel disease causing variants. We also searched for IBD across groups – screening for cases in which the across-group IBD score is at least a third of the within-group IBD score of Ashkenazi Jews – and found 77 cases of clear IBD-sharing, which typically follow geography, religious affiliation, or linguistic grouping (particularly Austroasiatic speakers) (Supplementary Table 4). Pairs of groups with high shared IBD and descent from a common founder event will share risk for the same recessive disease. However, these cross-group IBD sharing patterns are not driving our observations, as we still identify 68 unique sets of groups without high IBD to other groups that have significantly higher estimated IBD scores than both Finns and Ashkenazi Jews.

**Figure 3.**
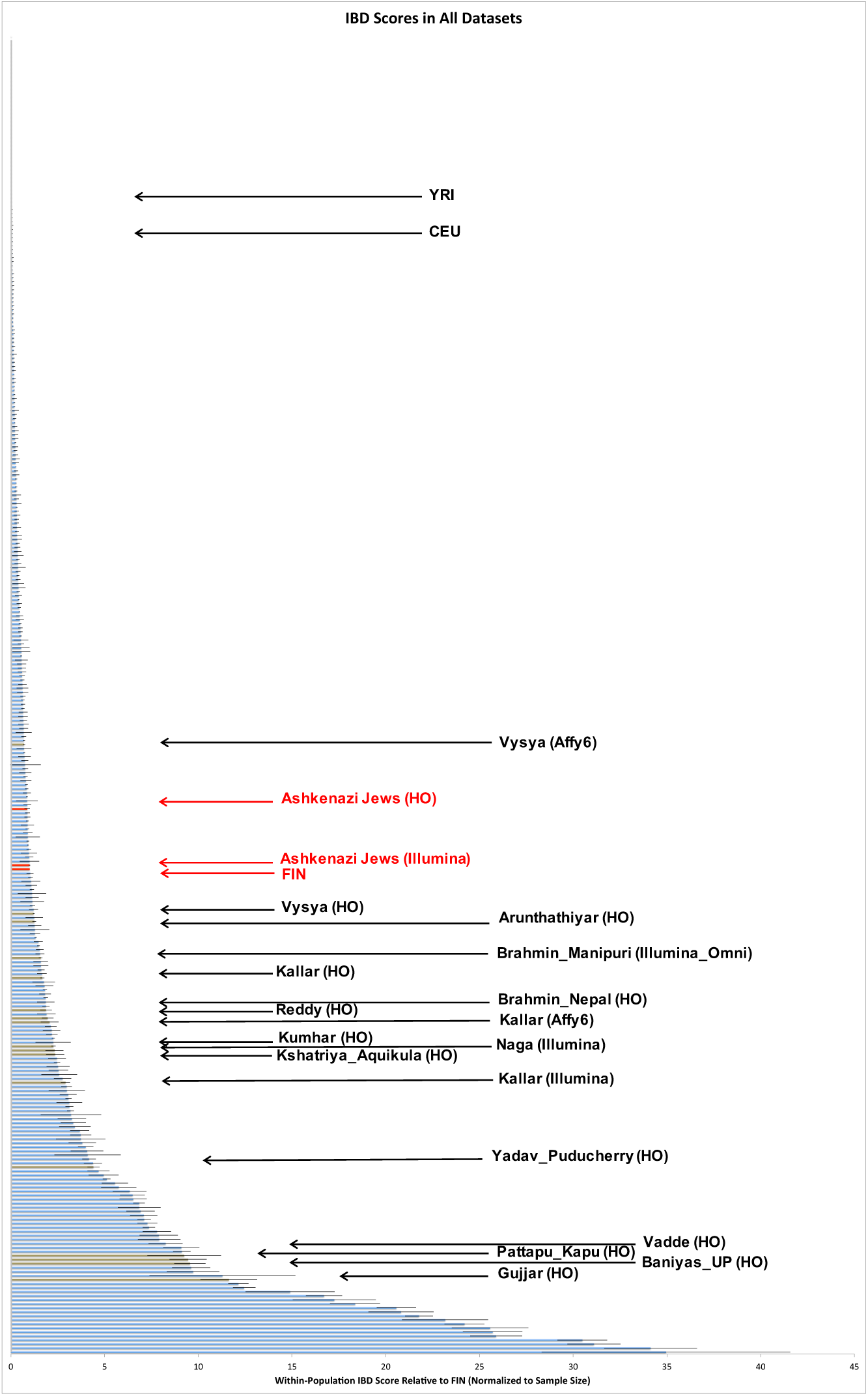
IBD scores relative to Finns (FIN). Histogram ordered by IBD score, roughly proportional to the per-individual risk for recessive disease due to the founder event. (These results are also given quantitatively for each group in Online Table 1.) We restrict to groups with at least two samples, combining data from all four genotyping platforms onto one plot. Data from Ashkenazi Jews and Finns are highlighted in red, and from South Asian groups with significantly higher IBD scores than that of Finns and census sizes of more than a million in brown. Error bars for each IBD score are standard errors calculated by weighted block jackknife over each chromosome. YRI = Yoruba (West African); CEU = European American.

**Table 1.** South Asian groups with estimated census sizes over 1 million and IBD scores significantly greater than those of Ashkenazi Jews and Finns. Fourteen South Asian groups with IBD scores significantly higher than that of Finns, census sizes over 1 million, and sample sizes of at least 3 that are of particularly high interest for founder event disease gene mapping studies. For reference, Finns and Ashkenazi Jews (on the Human Origins array) would have IBD scores of 1.0 and 0.9, IBD ranks of 121 and 135, and F_ST_ ranks of 109 and 129, respectively (the group-specific drift is difficult to compare for groups with significantly different histories, so they were not calculated for Finns or Ashkenazi Jews).

| Group | Sample Size | IBD Score | IBD Rank | F<sub>ST</sub> Rank | Drift Rank | Census Size | Location |
| --- | --- | --- | --- | --- | --- | --- | --- |
| Gujjar | 5 | 11.6 | 19 | 33 | 46 | 1,078,719 | Jammu and Kashmir |
| Baniyas | 7 | 9.6 | 24 | 22 | 18 | 4,200,000 | Uttar Pradesh |
| Pattapu_Kapu | 4 | 9.5 | 25 | 24 | 21 | 13,697,000 | Andhra Pradesh |
| Vadde | 3 | 9.2 | 26 | 30 | 26 | 3,695,000 | Andhra Pradesh |
| Yadav | 12 | 4.4 | 48 | 87 | 67 | 1,124,864 | Puducherry |
| Kshatriya_Aqnikula | 4 | 2.4 | 75 | 109 | NA | 12,809,000 | Andhra Pradesh |
| Naga | 4 | 2.3 | 76 | NA | NA | 1,834,483 | Nagaland |
| Kumhar | 27 | 2.3 | 77 | 35 | 197 | 3,144,000 | Uttar Pradesh |
| Reddy | 7 | 2.0 | 84 | 129 | 106 | 22,500,000 | Telangana |
| Brahmin_Nepal | 4 | 1.9 | 86 | 63 | 141 | 4,206,235 | Nepal |
| Kallar | 27 | 1.7 | 94 | 87 | 73 | 2,426,929 | Tamil Nadu |
| Brahmin_Manipuri | 17 | 1.6 | 99 | NA | NA | 1,544,296 | Manipur |
| Arunthathiyar | 18 | 1.3 | 108 | 109 | 81 | 1,192,578 | Tamil Nadu |
| Vysya | 39 | 1.2 | 110 | 46 | 35 | 3,200,000 | Telangana |

Our documentation that very strong founder events affect a large fraction of South Asian groups presents an opportunity for decreasing disease burden in South Asia. This source of risk for recessive diseases is very different from that due to marriages among close relatives, which is also a major cause of recessive disease in South Asia. To determine the relative impact of these factors, we computed FST, a measurement of allele frequency differentiation, between each group in the dataset and a pool of other South Asian groups chosen to be closest in terms of ancestry proportions. We find that inbreeding is not driving many of these signals, as 89 unique groups have higher FST scores than those of Ashkenazi Jews and Finns even after reducing the FST score by the proportion of allele frequency differentiation due to inbreeding. These results show that while most recessive disease gene mapping studies in South Asia have focused on families that are the products of marriages between close relatives, recessive diseases are also likely to occur at an elevated rate even in non-consanguineous cases because of shared ancestors more distantly in time.

As an example of the promise of founder event disease gene mapping in South Asia, we highlight the case of the Vysya, who have a census size of more than 3 million and who we estimate have an IBD score about 1.2-fold higher than Finns (Figure 3). The Vysya have an approximately 100-fold higher rate of butyrylcholinesterase deficiency than other groups, and Vysya ancestry is a known counter-indication for the use of muscle relaxants such as succinylcholine or mivacurium that are given prior to surgery^13^. This disease is likely to occur at a higher rate due to the founder event in Vysya’s history, and we expect that, like Finns, Vysya likely have a higher rate of many other diseases compared to other groups. Other examples of recessive disease genes with a likely origin in founder events are known anecdotally in South Asia, highlighting the importance of systematic studies to find them^14^.

To demonstrate how a new recessive disease in a founder event group could be mapped, we carried out whole genome SNP genotyping in 12 patients from southern India who had progressive pseudorheumatoid dysplasia (PPD), a disease known to be caused by mutations in the gene *WISP3^15,16^*. Of the 6 individuals with the Cys78Tyr mutation in *WISP3,*^15,16^ 5 were from non-consanguineous marriages, and we found a much higher fraction of IBD at the disease mutation site than in the rest of the genome in these individuals (Supplementary Figure 3a; Supplementary Figure 4a), consistent with the Cys78Tyr mutation that causes PPD in these patients owing its origin to a founder event. This pattern contrasts with the 6 other patients with different disease variants and 6 patients with a mutation causing a different disease (mucopolysaccharidosis (MPS) type IVA) who were from primarily consanguineous marriages, and who lacked significant IBD across their disease mutation sites, implying that in the case of these groups the driver for the recessive diseases was marriage between close relatives (Supplementary Text). This example highlights how not only marriages of close relatives, but also founder events, are substantial causes of rare recessive disease in South Asia.

The evidence of widespread strong founder events presents a major opportunity for disease gene discovery and public health intervention in South Asia that is not widely appreciated. The current paradigm for recessive disease gene mapping in South Asia is to collect cases in tertiary medical centers and map diseases in individuals with the same phenotype, often blinded to information about group affiliation as in the case of the PPD study where we do not have access to the ethnic group information. However, our results suggest that collecting information on group affiliation could be greatly strengthen the power of these studies. A fruitful way to approach gene mapping would be to proactively survey communities known to have strong founder events, searching for diseases that occur at high rates in these communities. This approach was pioneered in the 1950s in studies of the Old Order Amish in the U.S., a founder population of approximately 100,000 individuals in whom many dozens of recessive diseases were mapped, a research program that was crucial to founding modern medical genetics and that was of extraordinary health benefit. Our study suggests that the potential for disease gene mapping in South Asia would be orders of magnitude greater.

Mapping of recessive diseases may be particularly important in communities practicing arranged marriages, which are common in South Asia. An example of the power of this approach is given by *Dor Yeshorim*, a community genetic testing program among religious Ashkenazi Jews^17^, which visits schools, screens students for common recessive disease causing mutations previously identified to be segregating at a higher frequency in the target group, and enters the results into a confidential database. Matchmakers query the database prior to making suggestions to the families and receive feedback about whether the potential couple is “incompatible” in the sense of both being carriers for a recessive mutation at the same gene. Given that approximately 95% of community members whose marriages are arranged participate in this program, recessive diseases like Tay-Sachs have virtually disappeared in these communities. A similar approach should work as well in South Asian communities. Given the potential for saving lives, this or similar kinds of research could be a valuable investment for future generations^18^.

## Supplementary Data

Supplementary Data include an Excel spreadsheet detailing all groups and their scores on the IBD, F_ST_, and group-specific drift analyses. Also included are 7 supplementary figures and 5 supplementary tables.

## Acknowledgements

We are thankful to the many Indian, Pakistani, Bangladeshi, Sri Lankan, and Nepalese individuals who contributed the DNA samples analyzed here including the PPD and MPS patients. We are grateful to Analabha Basu and Partha P. Majumder for early sharing of data. Funding was provided by an NIGMS (GM007753) fellowship to NN, a Translational Seed Fund grant from the Dean’s Office of Harvard Medical School and an HG006399 to DR, Council of Scientific and Industrial Research, Government of India grant to KT, and an NIGMS grant 115006 to PM. SS and SMJ acknowledge the funding from the Department of Biotechnology (BT/PR4224/MED/97/60/2011) and Department of Science and Technology (SR/WOS-A/LS-83/2011), Government of India. Funding for the mutation analysis of Indian patients with PPD was provided by Indian Council of Medical Research (BMS 54/2/2013) to KMG. DR is an Investigator of the Howard Hughes Medical Institute. The informed consents and permits associated with the newly reported data are not consistent with fully public release. Therefore, researchers who wish to analyze the data should send the corresponding authors a PDF of a signed letter containing the following language: “(a) We will not distribute the data outside my collaboration, (b) We will not post data publicly, (c) We will make no attempt to connect the genetic data to personal identifiers, (d) We will not use the data for commercial purposes.”

## Author Contributions

N.N., P.M., D.R., and K.T. conceived the study. N.N., P.M., N.R., B.S., A.T., N.P. and D.R. performed analysis. G.B., K.M.G., M.S.M., S.S. A.K., S.A.V., S.M.J., K.S., L.S. and K.T. collected data. N.N., D.R., and K.T. wrote the manuscript with the help of all co-authors.

## Competing Financial Interests

The authors declare no competing financial interests.

Reprints and permissions information is available online at http://www.nature.com/reprints/index.html.

## Online Methods

### Data Sets

We assembled a dataset of 1,955 individuals from 249 groups genotyped on the Affymetrix Human Origins array, of which data from 1,663 individuals from 230 groups are newly reported here (Figure 1a). We merged these data with the dataset published in Moorjani *et al*.^8^, which consisted of 332 individuals from 52 groups genotyped on the Affymetrix 6.0 array. We also merged it with two additional datasets published in Metspalu *et al*.^9^, consisting of 151 individuals from 21 groups genotyped on Illumina 650K arrays as well as a dataset published in Basu *et al*.^11^, consisting of 367 individuals from 20 groups generated on Illumina Omni 1-Quad arrays. These groups come from India, Pakistan, Nepal, Sri Lanka, and Bangladesh. All samples were collected under the supervision of ethical review boards in India with informed consent obtained from all subjects.

We analyzed two different Ashkenazi Jewish datasets, one consisting of 21 individuals genotyped on Illumina 610K and 660K bead arrays^10^ and one consisting of 7 individuals genotyped on Affymetrix Human Origins arrays.

Our “Affymetrix 6.0” dataset consists of 332 individuals genotyped on 329,261 SNPs, and our “Illumina_Omni” dataset consists of 367 individuals genotyped on 750,919 SNPs. We merged the South Asian and Ashkenazi Jewish data generated by the other Illumina arrays to create an “Illumina” dataset consisting of 172 individuals genotyped on 500,640 SNPs. We merged the data from the Affymetrix Human Origins arrays with the Ashkenazi Jewish data and data from the Simons Genome Diversity Project^19,20^ to create a dataset with 4,402 individuals genotyped on 512,615 SNPs. We analyzed the four datasets separately due to the small intersection of SNPs between them. We merged in the 1000 Genomes Phase 3 data^21^ (2,504 individuals from 26 different groups; notably, including 99 Finnish individuals) into all of the datasets. We used genome reference sequence coordinates (hg19) for analyses.

### Quality Control

We filtered the data at both the SNP and individual level. On the SNP level, we required at least 95% genotyping completeness for each SNP (across all individuals). On the individual level, we required at least 95% genotyping completeness for each individual (across all SNPs).

To test for batch effects due to samples from the same group being genotyped on different array plates, we studied instances where samples from the same group *A* were genotyped on both plates 1 and 2 and computed an allele frequency difference at each SNP, 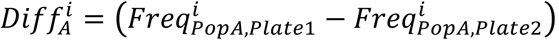. We then computed the product of these allele frequencies averaged over all SNPs for two groups A and B genotyped on the same plates, 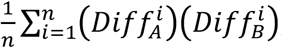, as well as a standard error from a weighted Block Jackknife across chromosomes. This quantity should be consistent with zero within a few standard errors if there are no batch effects that cause systematic differences across the plates, as allele frequency differences between two samples of the same group should be random fluctuations that have nothing to do with the array plates on which they are genotyped. This analysis found strong batch effects associated with one array plate, and we removed these samples from further analysis.

We used EIGENSOFT 5.0.1 smartpca^22^ on each group to detect PCA outliers and removed 51 samples. We also developed a procedure to distinguish recent relatedness from founder events so that we could remove recently related individuals. We first identified all duplicates or obvious close relatives by using Plink^23^ “genome” and GERMLINE^24^ to compute IBD (described in more detail below) and removed one individual from all pairs with a PI_HAT score greater than 0.45 and the presence of at least 1 IBD fragment greater than 30cM. We then used an iterative procedure to identify additional recently related individuals. For sample sizes above 5, we identified any pairs within each group that had both total IBD and total long IBD (>20cM) that were greater than 2.5 SDs and 1 SD, respectively, from the group mean. For sample sizes 5 or below, we used modified Z scores of 0.6745*(IBD_score - median(score))/MAD, where MAD is the median absolute deviation, and identified all pairs with modified Z scores greater than 3.5 for both total IBD and total long IBD as suggested by Iglewicz and Hoaglin^25^. After each round, we repeated the process if the new IBD score was at least 30% lower than the prior IBD score. Simulations showed that we were always able to remove a first or second cousin in the dataset using this method (Supplementary Table 1). Together these analyses removed 53 individuals from the Affymetrix 6.0 dataset, 21 individuals from the Illumina dataset, 43 individuals from the Illumina Omni dataset, and 225 individuals from the Human Origins dataset.

After data quality control and merging with the 1000 Genomes Project data, the Affymetrix 6.0 dataset included 2,842 individuals genotyped on 326,181 SNPs, the Illumina dataset included 2,662 individuals genotyped on 484,293 SNPs, the Illumina Omni dataset included 2,828 individuals genotyped on 750,919 SNPs, and the Human Origins dataset included 4,177 individuals genotyped at 499,158 SNPs.

### Simulations to Test Relatedness Filtering and IBD Analyses

We used ARGON^26^ to simulate groups with different bottleneck strengths to test the IBD analyses and relatedness filtering. We used ARGON’s default settings, including a mutation rate of 1.65*10^−8^ per base pair (bp) per generation and a recombination rate of 1*10^−8^ per bp per generation and simulated 22 chromosomes of size 130 Mb each. We pruned the output by randomly removing SNPs until there were 22,730 SNPs per chromosome to simulate the approximate number of positions in the Affymetrix Human Origins array. For the IBD analyses, we simulated groups to have descended from an ancestral group 1,800 years ago with N_e_ = 50,000 and to have formed two groups with N_e_ = 25,000. These groups continued separately until 100 generations ago when they combined in equal proportions to form a group with N_e_ = 50,000. The group then split into 3 separate groups 72 generations ago that have bottlenecks leading to Ne of either 400, 800, or 1600. The 3 groups then exponentially expanded to a present size of N_e_ = 50,000. We designed these simulations to capture important features of demographic history typical of Indian groups^4,8^. We chose the bottleneck sizes because they represent founder events with approximately the strength of Finns (the bottleneck to 800), and twice as strong (400) and half as strong (1600) as that group. We then performed the IBD analyses described below with 99 individuals from the group with bottleneck strength similar to that of Finns (198 haploid individuals were simulated and merged to produce 99 diploid individuals) and different numbers of individuals from the other groups. These analyses demonstrate that with only 4-5 individuals we can accurately assess the strength of founder events in groups with strong founder events (Supplementary Figure 2 and Supplementary Table 2). Weaker founder events are more difficult to assess, but these groups are of less interest for founder event disease mapping, so we aimed to sample ∼5 individuals per group.

We wrote custom R scripts to simulate first and second cousin pairs. We took individuals from the bottleneck of size 800 and performed “matings” by taking 2 individuals and recombining their haploid chromosomes assuming a rate of 1*10^−8^ per bp per generation across the chromosome and combining one chromosome from each of these individuals to form a new diploid offspring. The matings were performed to achieve first and second cousins. We then placed these back into the group with group of size 800, and ran the relatedness filtering algorithms to evaluate whether they would identify these individuals.

### Phasing, IBD Detection, and IBD Score Algorithm

We phased all datasets using Beagle 3.3.2 with the settings *missing=0; lowmem=true; gprobs=false; verbose=true*^27^. We left all other settings at default. We determined IBD segments using GERMLINE^24^ with the parameters *-bits 75 -err_hom 0 -err_het 0 -min_m 3*. We used the genotype extension mode to minimize the effect of any possible phasing heterogeneity amongst the different groups and used the HaploScore algorithm to remove false positive IBD fragments with the recommended genotype error and switch error parameters of 0.0075 and 0.003^28^. We chose a HaploScore threshold matrix based on calculations from Durand *et al.*^28^ for a “mean overlap” of 0.8, which corresponds to a precision of approximately 0.9 for all genetic lengths from 2-10cM. It can sometimes be difficult to measure IBD in admixed populations due to differential proportions of the divergent ancestries amongst different individuals in the same group, but we found that in both the simulated and real data we were able to detect IBD at the expected amounts.

In addition to the procedure we developed to remove close relatives (Quality Control section), we also removed segments longer than 20cM as simulations showed that this increased sensitivity of the analyses (Supplementary Table 2). We computed “IBD score” as the total length of IBD segments between 3-20cM divided by 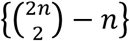 where n is the number of individuals in each group to normalize for sample size. We then expressed each group’s score as a ratio of their IBD score to that of Finns and calculated standard errors for this score using a weighted Block Jackknife over each chromosome with 95% confidence intervals defined as IBD score ±1.96*s.e.

We repeated these analyses with FastIBD^29^ for the Affymetrix 6.0 and Illumina datasets and observed that the results were highly correlated (r>0.96) (data not shown). We chose GERMLINE for our main analyses, however, because the FastIBD algorithm required us to split the datasets into different groups, since it adapts to the relationships between LD and genetic distance in the data, and these relationships differ across groups. We used data from several different Jewish groups and all twenty-six 1000 Genomes groups to improve phasing, but of these groups we only included results for Ashkenazi Jews and two outbred groups (CEU and YRI) in the final IBD score ranking.

### Disease patient analyses

We use Affymetrix Human Origins arrays to successfully genotype 12 patients with progressive pseudorheumatoid dysplasia (PPD) and 6 patients with mucopolysaccharidosis (MPS) type IVA, all of whom had disease mutations previously determined^15,16,30^ (3 of the surveyed MPS patients are newly reported here). A total of 6 of the PPD patients had Cys78Tyr mutations, 6 had Cys337Tyr mutations (all 6 of the MPS patients had Cys78Arg mutations). We measured IBD as described above and also detected homozygous segments within each individual by using GERMLINE with the parameters *-bits 75 -err_hom 2 -err_het 0 -min_m 0.5 -homoz-only*.

Haplotype sharing was assessed by analyzing phased genotypes for each mutation group. At each SNP, we counted the number of identical genotypes for each allele and computed the fraction by dividing by the total number of possible haplotypes (2 times the number of individuals). We took the larger value of the two possible alleles (thus the fraction range was 0.5-1). We averaged these values over blocks of 10 or 25 SNPs and plotted the averages around the relevant mutation site.

### Between-Group IBD Calculations

We determined IBD using GERMLINE as above. We collapsed individuals into respective groups and normalized for between-group IBD by dividing all IBD from each group by 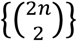 where n is the number of individuals in each group. We normalized for within-group IBD as described above. We defined groups with high shared IBD as those with an IBD score greater than three times the founder event strength of CEU (and ∼1/3 the event strength of Ashkenazi Jews).

### *f_3_*-statistics

We used the *f_3_*-statistic^7^ *f_3_(Test; Ref_1_, Ref_2_)* to determine if there was evidence that the *Test* group was derived from admixture of groups related to *Ref_1_* and *Ref_2_*. A significantly negative statistic provides unambiguous evidence of mixture in the Test group. We determined the significance of the *f_3_*-statistic using a Block Jackknife and a block size of 5 cM. We considered statistics over 3 standard errors below zero to be significant.

### Computing Group Specific Drift

We used qpGraph^7^ to model each Indian group on the cline as a mixture of ANI and ASI ancestry, using the model (YRI, (Indian group, (Georgians, ANI)), [(ASI, Onge])) proposed by Moorjani *et al*.^8^ This approach provides estimates for post-admixture drift in each group (Supplementary Figure 5), which is reflective of the strength of the founder event (high drift values imply stronger founder events). We only included groups on the Indian cline in this analysis, and we removed all groups with evidence of East Asian related admixture (Figure 1b and Supplementary Table 5) because this admixture is not accommodated within the above model.

### PCA-Normalized F_ST_ Calculations

As a third method to measure strength of founder events, we estimated the minimum F_ST_ between each South Asian group (Supplementary Figure 6) and their closest clusters based on PCA (Supplementary Text) (the clusters were used to account for intermarriage across groups that would otherwise produce a downward bias in the minimum F_ST_). For the Affymetrix 6.0, Illumina, and Illumina_Omni datasets, we split the Indian cline into two different clusters and combined the Austroasiatic speakers and those with ancestry related to Austroasiatic speakers (according to the PCA of Figure 1b) into one cluster for a total of three clusters (all other groups were ignored for this analysis). For the Human Origins dataset we split the Indian cline into three different clusters and combined the groups with ancestry related to the main cluster of Austroasiatic speakers into one cluster for a total of four clusters (Khasi and Nicobarese were ignored in this analysis, because they do not cluster with the other Austroasiatic speaking groups). We then computed the F_ST_ between each group and the rest of the individuals in their respective cluster based on EIGENSOFT *smartpca* with Inbreed set to YES to correct for inbreeding. For Ashkenazi Jews and Finns, we used the minimum F_ST_ to other European groups.

### F_ST_ Calculations to Determine Overlapping Groups

Overlapping groups between the datasets were determined in the first place based on anthropological information (Online Table 1). We further tested empirically for overlap by computing F_ST_ between different groups across all datasets for groups with significantly stronger IBD scores than those of Finns (we could not perform this analysis for groups with less strong founder events, because they would have low F_ST_ to each other even if they were truly distinct groups). We considered pairs with F_ST_ less than 0.004 to be overlapping. These included all groups known to be overlapping based on anthropological information as well as 3 additional pairs of groups that might be genetically similar due to recent mixing (e.g. Kanjars and Dharkar are distinct nomadic groups that live near each other but intermarry, leading to low F_ST_ between them).

### Code Availability

Code for all calculations available upon request.

